# Deficiency of peroxisomal L-bifunctional protein (EHHADH) causes male-specific kidney hypertrophy and proximal tubular injury in mice

**DOI:** 10.1101/2021.03.14.435187

**Authors:** Pablo Ranea-Robles, Kensey Portman, Aaron Bender, Kyung Lee, John Cijiang He, David J Mulholland, Carmen Argmann, Sander M Houten

**Affiliations:** Department of Genetics and Genomic Sciences, Icahn Institute for Data Science and Genomic Technology, Icahn School of Medicine at Mount Sinai, New York, NY 10029, USA; Division of Oncological Sciences, Icahn School of Medicine at Mount Sinai, New York, NY 10029, USA; Department of Medicine, Division of Nephrology, Icahn School of Medicine at Mount Sinai, New York, NY 10029, USA

**Keywords:** Peroxisomal bifunctional protein, Multifunctional protein 1, kidney, sex differences, acute kidney injury

## Abstract

**Background:** Proximal tubular (PT) cells are enriched in mitochondria and peroxisomes. Whereas mitochondrial fatty acid oxidation (FAO) plays an important role in kidney function by supporting the high-energy requirements of PT cells, the role of peroxisomal metabolism remains largely unknown. EHHADH, also known as L-bifunctional protein, catalyzes the second and third step of peroxisomal FAO.

**Methods:** We studied kidneys of WT and *Ehhadh* KO mice using histology, immunohistochemistry, immunofluorescence, immunoblot, RNA-sequencing, metabolomics and orchiectomy.

**Results:** We observed male-specific kidney hypertrophy and glomerular filtration rate reduction in adult *Ehhadh* KO mice. Transcriptome analysis unveiled a gene expression signature similar to PT injury in acute kidney injury mouse models. This was further illustrated by the presence of KIM-1 (kidney injury molecule-1), SOX-9, and Ki67-positive cells in the PT of male *Ehhadh* KO kidneys. Male *Ehhadh* KO kidneys had metabolite changes consistent with peroxisomal dysfunction as well as an elevation in glycosphingolipids levels. Orchiectomy of *Ehhadh* KO mice reversed kidney enlargement and decreased the number of KIM-1 positive cells. We reveal a pronounced sexual dimorphism in the expression of peroxisomal FAO proteins in mouse kidney, underlining a role of androgens in the kidney phenotype of *Ehhadh* KO mice.

**Conclusions:** Our data highlight the importance of EHHADH and peroxisomal metabolism in male kidney physiology and reveal peroxisomal FAO as a sexual dimorphic metabolic pathway in mouse kidneys.

## Introduction

The kidney uses the mitochondrial fatty acid β-oxidation (FAO) pathway as the predominant energy source (Nieth & Schollmeyer 1966; Weidemann & Krebs 1969) to fulfill the high energy requirements of proximal tubule (PT) cells (Kang et al. 2015). Dysfunction of mitochondrial FAO has been linked to the development of kidney fibrosis in chronic kidney disease (CKD) patients and mouse models (Afshinnia et al. 2018; Dhillon et al. 2021; Kang et al. 2015; Miguel et al. 2021). PT cells are not only enriched in mitochondria, but also in peroxisomes (Rhodin 1954). Peroxisomes have unique metabolic functions that include the β-oxidation of specific carboxylic acids such as very long-chain fatty acids, and the biosynthesis of plasmalogens (Wanders & Waterham 2006). The importance of peroxisomes in kidney function is highlighted by the presence of renal cysts and/or calcium oxalate stones in patients with Zellweger spectrum disorder (ZSD) and other peroxisomal diseases (Goldfischer et al. 1973; Huyghe et al. 2006; Steinberg et al. 2006; van Woerden et al. 2006). Moreover, peroxisome abundance and function are reduced in several rodent models of kidney injury (Gulati et al. 1992; Kalakeche et al. 2011; Malas et al. 2017; Negishi et al. 2007; Ruidera et al. 1988). These data indicate that peroxisomes are important for kidney function. The exact roles of peroxisomes in the kidney, however, remain unknown.

Each cycle of peroxisomal FAO consists of four enzymatic steps. The second and third steps are catalyzed by the peroxisomal L- and D-bifunctional proteins (encoded by *EHHADH* and *HSD17B4*, respectively). EHHADH is mainly expressed in liver and kidney. We and others have characterized a specific role of EHHADH in the hepatic metabolism of long-chain dicarboxylic acids (DCAs) (Ding et al. 2013; Ferdinandusse et al. 2004; Houten et al. 2012; Nguyen et al. 2008; Ranea-Robles et al. 2021), but the role of EHHADH in the metabolic homeostasis of the kidney is currently unknown.

In the present study, we examined the role of EHHADH in the kidney by using a KO mouse model (*Ehhadh* KO mice). Our results demonstrate that the *Ehhadh* KO mouse is a new model for metabolic kidney injury with enhanced susceptibility in male mice, and underline the role of peroxisomes in kidney physiology.

## Methods

### Animal experiments

All animal experiments were approved by the IACUC of the Icahn School of Medicine at Mount Sinai (# IACUC-2014-0100) and comply with the National Institutes of Health guide for the care and use of Laboratory animals (NIH Publications No. 8023, 8th edition, 2011). The generation of *Ehhadh* KO (*Ehhadh*^-/-^) mice has been previously described (Qi et al. 1999; Violante et al. 2019). WT and *Ehhadh* KO mice on a pure C57BL/6N background were fed a regular chow (PicoLab Rodent Diet 20, LabDiet). Mice were housed in rooms with a 12 h light/dark cycle. Mice were euthanized by exposure to CO_2_ and blood was collected from the inferior vena cava for the preparation of EDTA plasma. Organs were snap frozen in liquid nitrogen and stored at −80°C.

### Orchiectomy

To remove the production of gonadal androgens, surgical castration surgery was carried out in WT and *Ehhadh* KO mice under full anesthesia, as previously described (Liang et al. 2015).

### Blood urea nitrogen and plasma creatinine

Measurements of blood urea nitrogen (BUN) and creatinine were performed in mouse plasma. BUN was measured using a quantitative colorimetric QuantiChrom kit (DIUR-100, BioAssay Systems), following manufacturer’s instructions. Creatinine was analyzed as described (Le et al. 2014).

### Glomerular filtration rate

Glomerular filtration rate (GFR) was measured in conscious mice using clearance kinetics of plasma FITC-inulin after a single bolus injection in the vein tail, as previously described (Qi et al. 2004). GFR was calculated based on a two-compartment model. GFR was expressed in µL per minute.

### Histology, immunohistochemistry and immunofluorescence

Kidneys were collected after CO_2_ euthanasia, weighed, cut in two transverse halves, and fixed by immersion in 10% formalin (Thermo Fisher Scientific) for 24 hours. Next, fixed kidneys were washed in PBS, transferred to 70% EtOH and embedded in paraffin blocks. Serial transversal sections (4 µm thick) were cut with a microtome. Immunohistochemistry (IHC) and immunofluorescence (IF) studies were carried out using the avidin-peroxidase method and fluorescent antibodies, respectively. For both IHC and IF, we performed antigen retrieval by boiling preparations 20 minutes in 5 mM citrate buffer pH 6.0. Next, kidney sections were blocked in 0.3% Triton X-100 in 5% donkey normal serum in PBS.

The incubation with primary antibody was performed in 0.1% Triton X-100 in 3% donkey normal serum in PBS, overnight at 4°C, using the following antibodies: anti-KIM-1 (AF1817, RD Systems), anti-SOX-9 (NBP1-8551, Novus Bio), anti-LRP2 (ab76969, Abcam), anti-SGLT1 (ab14685, Abcam), biotinylated-LTL (B-1325, Vector Laboratories), anti-EHHADH (GTX81126, Genetex), anti-Ki67 (MA5-14520, Invitrogen), anti-AQP2 (AQP-002, Alomone labs), anti-CALB (PA5-85669, Thermo Fisher), and anti-Slc34a3 (NPT2c, 20603, BiCell Scientific). The incubation with secondary antibody was performed in PBS for 2 hours at room temperature, using the following secondary antibodies: 705-585-147, 712-545-153, 016-540-084 (Jackson ImmunoResearch), and A21206 (Invitrogen).

When two primary antibodies raised in rabbit were required for double IF, sections were incubated with AffiniPure Fab Fragment Goat Anti-Rabbit IgG (111-007-003) after the first primary antibody incubation in order to allow its detection with anti-goat secondary antibody. Control slides confirmed that the first primary antibody was not detected by the anti-rabbit secondary antibody (data not shown). Nuclei were visualized with a Hoechst stain. For IF studies, autofluorescence was reduced by incubating the slides in 0.3% Sudan Black in 70% EtOH (Electron Microscopy Sciences) for 15 minutes. Microscopy images were taken with a Nikon Eclipse 80i microscope and the NIS-Elements BR 5.20.01 software (Nikon). Images were analyzed with ImageJ (Schindelin et al. 2012; Schneider et al. 2012). The number of Ki67^+^ cells per mm^2^ and the number of double SOX-9/KIM-1 positive cells in the kidney were determined by counting positive cells with the Cell Counter plugin in ImageJ (Schindelin et al. 2012; Schneider et al. 2012). The cross-sectional area (in µm^2^) of 50 random cortical tubules was measured in H&E-stained sections of WT and *Ehhadh* KO kidneys using the ROI manager tool of ImageJ (Schindelin et al. 2012; Schneider et al. 2012). The researcher was blinded to the genotype of the sample when analyzing the images.

### Immunoblot analysis

Protein was extracted from frozen mouse kidneys and immunoblot analysis was performed as previously described (Ranea-Robles et al. 2020) using the following primary antibodies: anti-KIM-1 (AF1817, RD Systems), anti-SOX-9 (NBP1-8551, Novus Bio), anti-SPTLC2 (51012-2-AP, Proteintech), anti-ABCD3 (PA1-650, Invitrogen), anti-ACOX1 (ab184032, Abcam), anti-EHHADH (GTX81126, Genetex), anti-HSD17B4 (15116-1-AP, Proteintech), anti-ACAA1 (12319-2-AP, Proteintech), anti-SCPx (HPA027135, Atlas Antibodies), anti-CROT (NBP1-85501, Novus Bio), anti-CPT2 (26555-1-AP, Proteintech), anti-MCAD (55210-1-AP, Proteintech), anti-α-tubulin (32-2500, Thermo Fisher), and anti-citrate synthase (GTX628143, Genetex). The anti-AMACR antibody was a kind gift from Dr. Sacha Ferdinandusse (AMC, Amsterdam) (Ferdinandusse et al. 2000).

### RNA-seq, differential gene expression, pathway enrichment

RNA was isolated using QIAzol lysis reagent followed by purification using the RNeasy kit (Qiagen). RNA samples were submitted to the Genomics Core Facility at the Icahn Institute and Department of Genetics and Genomic Sciences for further processing. Briefly, mRNA-focused cDNA libraries were generated using Illumina reagents (polyA capture), and samples were run on an Illumina HiSeq 2500 sequencer to yield a read depth of approximately 56 million 100 nucleotide single-end reads per samples. Reads from fastq files were aligned to the mouse genome mm10 (GRCm38.75) with STAR (release 2.4.0 g1) (Dobin et al. 2013) and summarized to gene- and exon-level counts using featureCounts (Liao et al. 2014). Only genes with at least one count per million in at least 2 samples were considered. Differential gene expression analysis was conducted with the R package limma (Law et al. 2014; Ritchie et al. 2015), as previously described (Argmann et al. 2017) Differentially expressed genes (DEGs) were defined using an adjusted p value of < 0.05 with no logFC cut-off.

Pathway enrichment analysis was performed using the Fisher’s exact test (FET) and p values were adjusted using Benjamini-Hochberg (BH) procedure. Hallmark and BioPlanet Pathways were sourced from Enrichr(Chen et al. 2013; Kuleshov et al. 2016). Input genes included genes up- or down-regulated (at adj p < 0.05) in *Ehhadh* KO vs WT kidneys that were converted from mouse to human orthologs using g:Orth from g:Profiler (Raudvere et al. 2019). iRegulon v1.3 was used to predict transcriptional regulators of kidney DEGs (Janky et al. 2014). Input genes included genes up- or down-regulated (at adj p < 0.05) in *Ehhadh* KO vs WT kidneys that were converted from mouse to human orthologs. The following options Motif collection: 10 K (9713 PWMs), Putative regulatory region: 10 kb centered around TSS (10 species) alongside the program default settings for Recovery and TF prediction options were selected for the analysis.

Genesets associated with murine PT responses to acute injury were curated from a clustering analysis of single cell RNAseq analysis of kidneys sampled over 12 h and 2, 14, and 42 days after bilateral –reperfusion (Kirita et al. 2020). Genesets associated with 8 sub-clusters, namely healthy S1, S2 and S3, repairing PT, injured S1/2, injured S3, severe injured PT and failed repair PT were curated from Dataset S2 of Kirita et al (Kirita et al. 2020). Genes found differentially expressed in FAN (folic acid nephropathy) versus control mice in PT cells were curated from Supplementary Table S2 of Dhillon et al (Dhillon et al. 2021). Genes found differentially expressed in PT after unilateral ureteral obstruction (UUO) versus control mice (at adj p < 0.01) were curated from Supplemental Table 2 of Wu et al (Wu et al. 2020). PT injury associated murine genesets were tested for enrichment in up- or down-regulated DEGs (at adj p < 0.05) between *Ehhadh* and WT mice using the FET and p values were adjusted using BH procedure.

Genes with sex-specific expression differences in mouse kidney were curated from two sources. One set were the gene differentially expressed (at adj p < 0.05) between kidneys of 10-week-old healthy BALB/c male and female mice (n=5 per group) (Si et al. 2009). The other set were genes differentially translated (at adj p < 0.05) in PT of contralateral kidneys from mice subjected to UUO (n=3 females, n=4 males) (Wu et al. 2020). DEGs were converted from mouse to human orthologs using g:Orth from g:Profiler (Raudvere et al. 2019). Pathway enrichment analysis was performed using Enrichr.(Chen et al. 2013; Kuleshov et al. 2016)

### Metabolomics

Global metabolite profiling (mView) from kidney (half, transversal) samples of 7 WT males and 5 *Ehhadh* KO males was performed by Metabolon, Inc. (Research Triangle Park, NC). To remove protein, to dissociate small molecules bound to protein or trapped in the precipitated protein matrix, and to recover chemically diverse metabolites, proteins were precipitated with methanol under vigorous shaking for 2 min (Glen Mills GenoGrinder 2000) followed by centrifugation. The resulting extract was analyzed by two separate reverse phase (RP)/UPLC-MS/MS methods with positive ion mode electrospray ionization (ESI), one RP/UPLC-MS/MS method with negative ion mode ESI, and one HILIC/UPLC-MS/MS method with negative ion mode ESI, as previously described (Miller et al. 2015). The scaled imputed data (Scaled Imp Data) represent the normalized raw area counts of each metabolite rescaled to set the median equal to 1. Any missing values were imputed with the minimum value. Metabolite pathway enrichment analysis using significantly altered HMDB metabolites was performed using MetaboAnalyst platform (Chong et al. 2018; Xia & Wishart 2011).

### Statistical analysis

Data are displayed as the mean ± the standard deviation (SD), with individual values shown. Differences were evaluated using unpaired t test with Welch’s correction or two-way ANOVA, as indicated in the figure legends. Significance is indicated in the figures. GraphPad Prism 8 was used to compute statistical values.

## Results

### EHHADH deficiency induces male-specific kidney hypertrophy without signs of severe pathology

We observed male-specific kidney enlargement in 4 to 9 month-old *Ehhadh* KO mice (**Fig. 1A**). Kidney mass and kidney-to-body weight (BW) ratio were higher in male *Ehhadh* KO mice compared to WT male mice (**Fig. 1B, C**). No changes in kidney mass were observed in female *Ehhadh* KO mice (**Fig. 1B, 1C**). Histological analysis revealed that *Ehhadh* KO mice showed male-specific proximal tubule (PT) hypertrophy (**Fig. 1D, 1E**, **S1A and S1B**), but changes suggestive of kidney damage were not observed (**Fig. 1D and S1A**). Glomerular filtration rate (GFR) was decreased in male *Ehhadh* KO mice when compared with WT mice (**Fig. 1F**), but this was not accompanied by alterations in circulating kidney function markers such as plasma creatinine (**Fig. 1G**) or blood urea nitrogen (BUN) (**Fig. 1H**). GFR and BUN in female *Ehhadh* KO mice were similar to WT **(Fig. S1C, S1D)**. In summary, EHHADH deficiency causes male-specific kidney hypertrophy and a decrease in GFR without signs of severe kidney damage.

**Figure 1.**
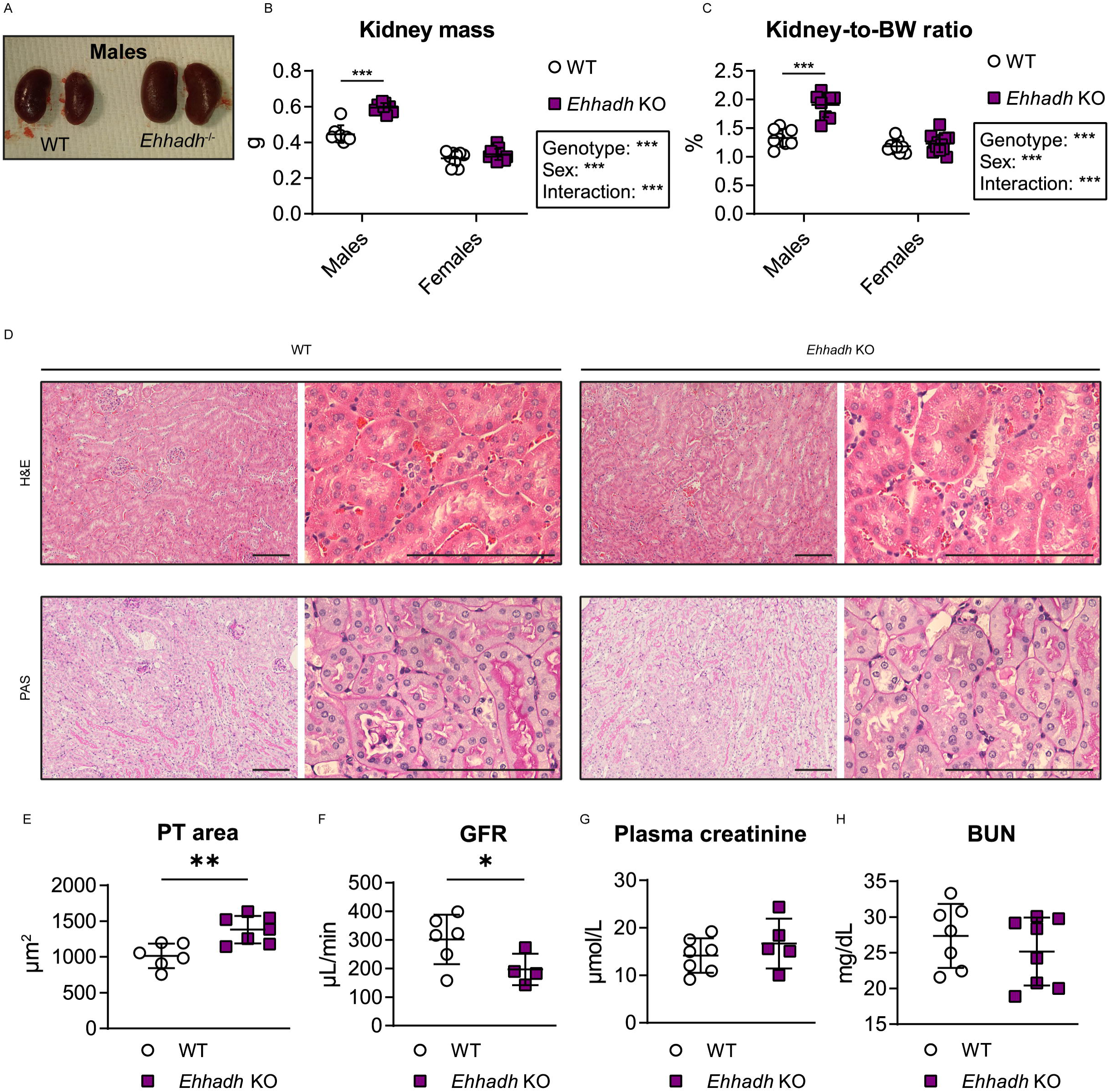
EHHADH deficiency induces male-specific kidney hypertrophy without signs of severe pathology. **A)** Representative images of WT and *Ehhadh* KO kidneys from male mice. **B, C**) Measurement of **B**) kidney weight (combined weight of both kidneys per animal) and **C**) kidney-to-body weight (BW) ratio in 4 to 9 month-old male WT (n=8), male *Ehhadh* KO (n=9), female WT (n=9), and female *Ehhadh* KO mice (n=12). **D**) Representative images of H&E and PAS staining of kidney sections from male WT and *Ehhadh* KO mice. Scale bars = 100 µm. **E**) Morphometric analysis of the cross-sectional tubule areas in WT (n=6) and *Ehhadh* KO (n=7) male mice **F**) Glomerular filtration rate (GFR) in WT (n=6) and Ehhadh KO (n=4) male mice. **G**) Plasma creatinine levels (μ in WT (n=5) and *Ehhadh* KO (n=7) male mice. **H**) Blood urea nitrogen (BUN) levels (mg/dL) in WT (n=7) and *Ehhadh* KO (n=8) mice male. Data are presented as mean ± SD with individual values plotted. ***P < 0.001, **P < 0.01, *P < 0.05, by two-way ANOVA (**B** and **C**), or unpaired t-test with Welch’s correction (**E, F**).

### Transcriptional activation of proximal tubule kidney injury signatures in male *Ehhadh* KO kidneys

We performed RNA-sequencing (RNA-seq) analysis on whole kidneys from male adult WT and *Ehhadh* KO mice (n=4). We identified 1475 significantly differentially expressed genes (DEGs, at adj p < 0.05), of which 806 genes were up- and 669 genes were down-regulated (**Fig. 2A and Table S1A**). Among the top up-regulated genes, we identified the genes encoding for kidney injury molecule 1 (KIM-1, encoded by *Havcr1*) and the transcription factor SOX-9 (*Sox9*), which are considered markers of PT injury (Han et al. 2002; Ichimura et al. 1998, 2004; Kuehn et al. 2002) and PT regeneration (Kang et al. 2016; Kumar et al. 2015), respectively (**Fig. 2A and Table S1A**).

**Figure 2.**
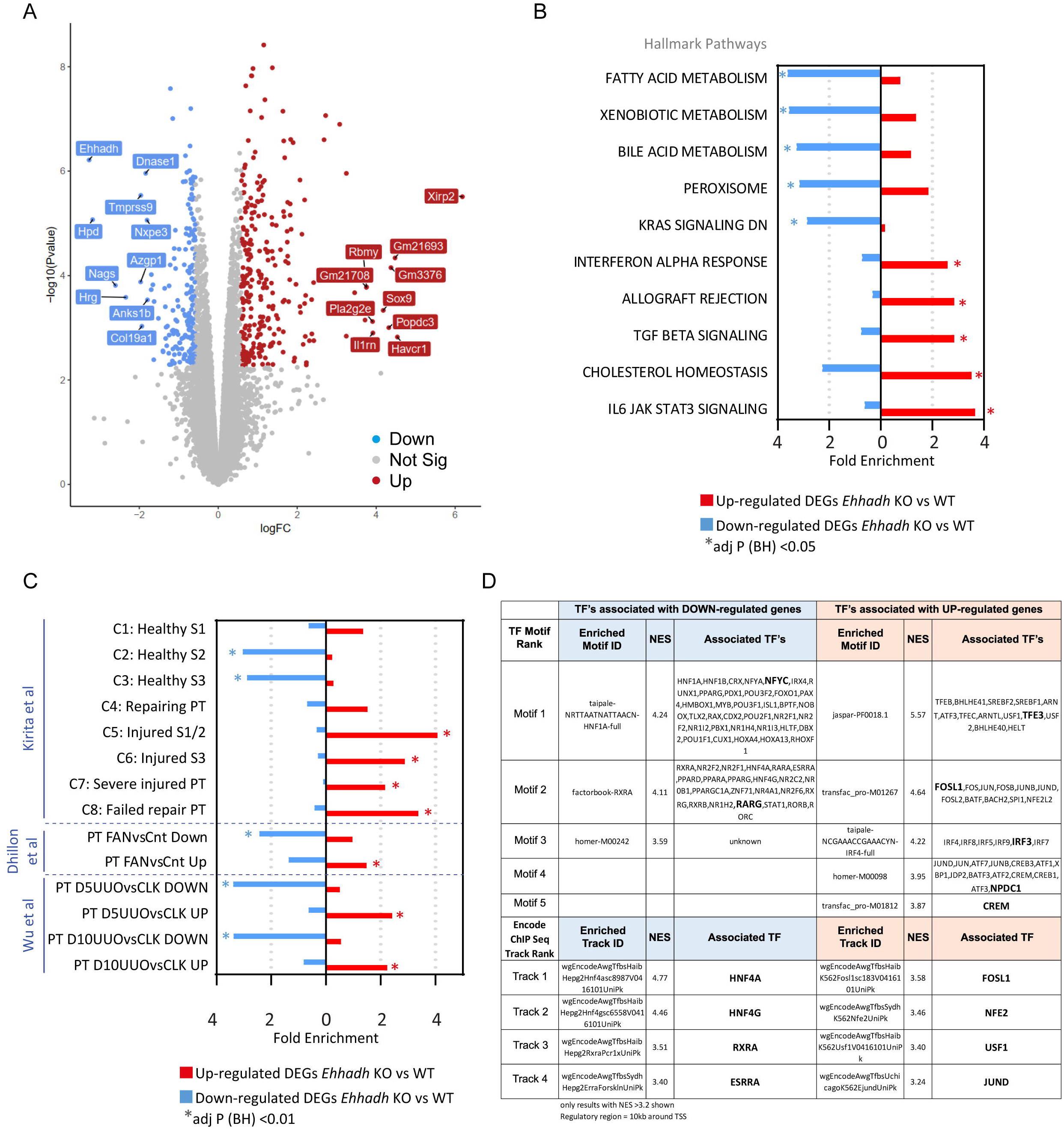
Transcriptomics of male *Ehhadh* KO kidneys. **A**) Volcano plot with significantly differentially expressed genes (DEGs) highlighted. Decreased genes (down) are highlighted in blue, increased genes (up) are highlighted in red. Gene names indicate the top 10 significant up and down genes by logFC using an adj p < 0.05. Genes that were not statistically significant are represented in grey. **B)** Top 10 pathways (by fold enrichment) after pathway enrichment analysis of genes either significantly up- or down-regulated in *Ehhadh* KO mice versus WT according to the Hallmark database. Values represent the fold enrichment and significance is indicated as * (adj p<0.05). Full table of results are in Table S1B and S1C. **C)** Results of enrichment analysis of genes either up- or down-regulated in *Ehhadh* KO mice versus WT, in gene sets curated from three independent murine proximal tubule (PT) acute kidney injury models (see Methods). Values represent the fold enrichment and significance is indicated as * (adj p<0.01). Full table of results are in Table S1D. **D)** Predicted transcriptional regulators as identified by Iregulon through Motif or Encode ChIP-seq enrichment analysis for the genes either up- or down-regulated in *Ehhadh* KO mice versus WT kidney samples. Only results with normalized enrichment score (NES) >3.2 are shown. Full table of results are in Table S1E. The bolded transcription factor (TF) represents the most likely TF associated with the enriched motif. Human orthologs of the murine DEGs (at adj p<0.05) were the input for B), C) and D).

Pathway enrichment analysis of the down-regulated DEGs using the Hallmarks database highlighted “Fatty acid metabolism”, “Bile acid metabolism”, and “Peroxisome”, among other pathways (**Fig. 2B, Fig. S2A, Table S1B**). Pathway enrichment analysis of the up-regulated DEGs highlighted inflammatory pathways, such as “Interferon alpha response”, “TGF beta signaling”, and “IL6 JAK STAT3 signaling”. Similar pathways were found enriched using the Bioplanet database (**Fig. S2B, Table S1C**). These changes resemble reported transcriptional changes in kidneys after acute kidney injury (AKI) (Dhillon et al. 2021; Kang et al. 2015; Kirita et al. 2020; Wu et al. 2020), suggesting that EHHADH deficiency causes kidney injury in male mouse kidneys.

We then compared the DEGs identified in *Ehhadh* KO male kidneys with expression signatures from PT cells of three different AKI mouse models; ischemia-reperfusion injury (Kirita et al. 2020), folic acid nephropathy (FAN) (Dhillon et al. 2021) and unilateral ureteral obstruction (UUO) (Wu et al. 2020) (**Table S1D**). Genes upregulated or downregulated in the male *Ehhadh* KO kidneys generally changed in the same direction in the FAN and UUO models (**Fig. 2C**). A recent report of Kirita et al. identified different cell states in the PT of mice subjected to ischemia-reperfusion injury, including a distinct proinflammatory and profibrotic PT cell state that fails to repair (Kirita et al. 2020). Our enrichment analysis revealed a significant enrichment of down-regulated DEGs from *Ehhadh* KO kidneys in the healthy S2 and healthy S3 PT cell states, and an enrichment of up-regulated DEGs in the injured S1/2, the injured S3, the severe injured PT and the failed repair PT cell states (**Fig. 2C**).

Next, we performed *cis*-regulatory sequence analysis to uncover the transcriptional regulatory network underlying the transcriptional response of *Ehhadh* KO male mouse kidneys (Janky et al. 2014) (**Table S1E**). We identified enriched transcription factor (TF) motifs in the down-regulated DEGs of *Ehhadh* KO kidneys that mapped to known regulators of PT differentiation such as HNF1A, HNF4A, RXRA, and ESRRA (Dhillon et al. 2021; Kirita et al. 2020; Marable et al. 2020) (**Fig. 2D**). In the up-regulated DEGs, we found enriched TF motifs that mapped to regulators of lysosomal biogenesis (TFEB/TFE3) (Raben & Puertollano 2016), and TFs whose activation is associated with PT cell injury (FOSL1) (Kirita et al. 2020), among others (**Fig. 2D**). In summary, these data show that the transcriptional response of male mouse kidney to EHHADH deficiency is very similar to the transcriptional signature present in AKI mouse models and is characteristic for PT cell injury.

### EHHADH deficiency activates the proximal tubule injury response in male mice

To validate the results of the RNA-seq, we performed immunohistochemistry (IHC) staining of KIM-1^+^ and identified KIM-1 cells, indicative of cellular injury (Ichimura et al. 2004), in male *Ehhadh* KO kidneys, in contrast to the rare appearance or absence of KIM-1^+^ cells in male WT kidneys and female WT and *Ehhadh* KO kidneys (**Fig. 3A**). KIM-1^+^ cells were scattered among tubules in *Ehhadh* KO male kidneys with a prominent cytoplasmic pattern, combined with some tubular cells that showed the typical apical location of KIM-1 that is present in acute kidney injury (Ichimura et al. 2004). We further validated the increase in KIM-1 protein in renal homogenates of male *Ehhadh* KO kidneys (**Fig. S3A**).

**Figure 3.**
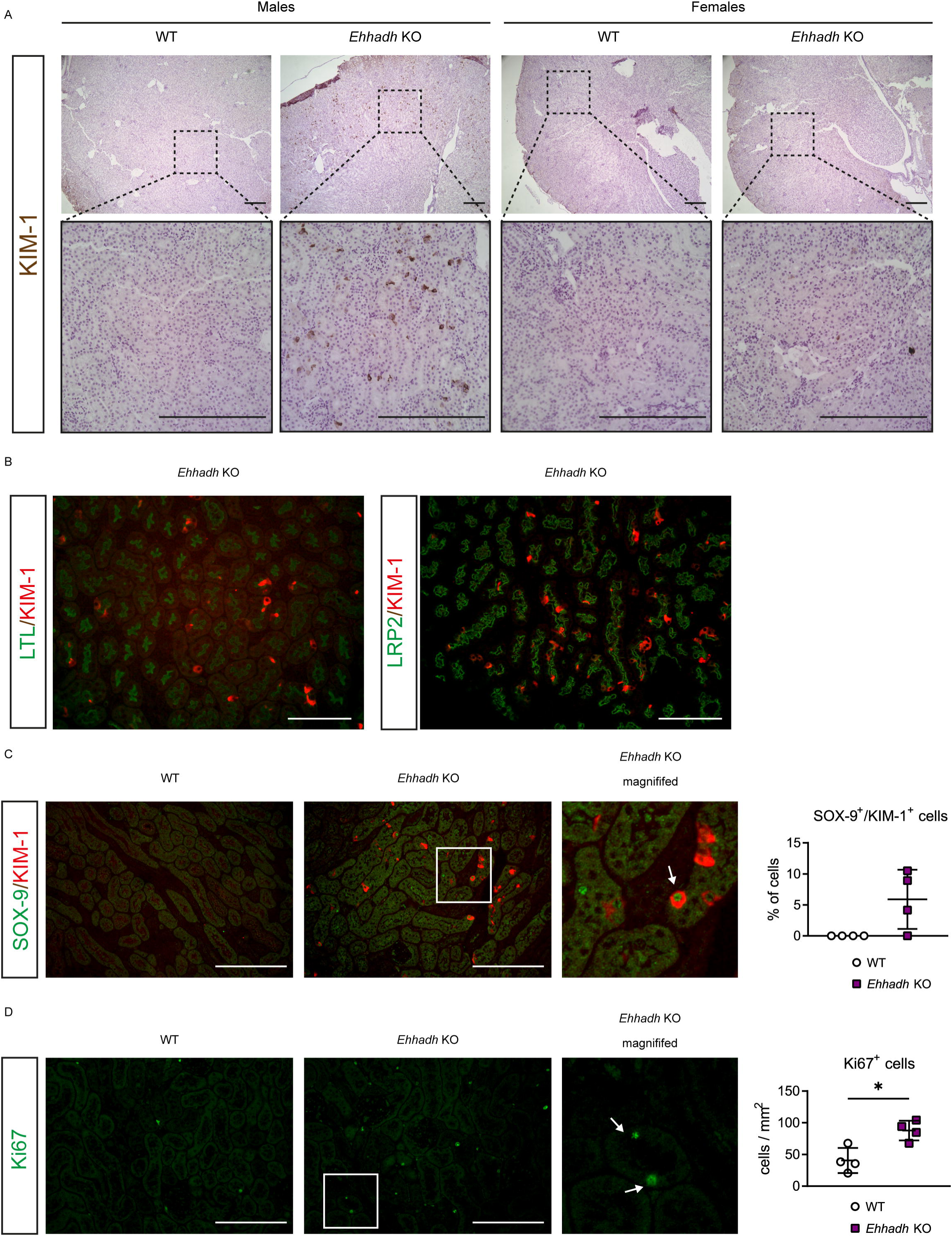
EHHADH deficiency activates the proximal tubule injury response in male mice. **A)** Representative images of KIM-1 IHC in the cortical area of WT and *Ehhadh* KO mouse kidneys (n=3-4 per sex and genotype). Upper panels: 4x objective. Lower panels: 20x objective. **B**) Representative images of double IF of KIM-1 with pan-tubular markers (LTL and LRP2) in the kidneys of *Ehhadh* KO male mice. There was no KIM-1 signal in male WT mice or female WT and *Ehhadh* KO mice (not shown). **C**) Representative images of double IF of KIM-1 (red) and SOX-9 (green) in WT and *Ehhadh* KO male mice (n=4). Inset shows expanded region of the *Ehhadh* KO section. White arrow points to a double positive Sox9^+^/Kim-1^+^ cell. Graph shows quantification of double positive SOX-9^+^/KIM-1^+^ cells in WT and *Ehhadh* KO mice. **D**) Representative images of Ki67 IF (green) in WT and *Ehhadh* KO male mice (n=4). Inset shows expanded region of the *Ehhadh* KO section. White arrows point to Ki67^+^ -cells. Graph shows quantification of Ki67^+^ cells per mm^2^ in WT and *Ehhadh* KO mice. Data are presented as mean ± SD with individual values plotted (**C, D**). Statistical significance was tested using unpaired t test with Welch’s correction (**C, D**). *P < 0.05. Scale bar = 250 µm (**A**), 100 µm (**B-D**).

Using double immunofluorescence (IF), we localized the KIM-1^+^ cells in the PT of *Ehhadh* KO male kidneys using the pan-tubular markers LTL and Megalin (LRP2) (**Fig. 3B**). Most of the KIM-1^+^ cells colocalized with SGLT1 (SLC5A1), a marker for the S2/S3 PT segments, whereas a small number of KIM-1^+^ cells colocalized with the S1 marker NPT2c (SLC34A3) (**Fig. S3B and S3C**). The localization of the majority of KIM-1^+^ cells in the S2/S3 segments is consistent with the expression pattern of EHHADH in mouse kidney, which is higher in S2/S3 than in S1 (Cheval et al. 2011; Limbutara et al. 2020) **(Fig S3D, S3E**). We did not find colocalization between KIM-1 and markers for distal tubule cells (CALB) or principal cells (AQP2) (**Fig. S3F, S3G**).

Double IF against KIM-1 and the transcription factor SOX-9 showed the presence of KIM-1^+^ and SOX-9^+^ cells in *Ehhadh* KO male kidneys (**Fig. 3C**). Among the KIM-1 cells, 5.9 ± 4.8% were also SOX-9^+^ (**Fig. 3C**), reflecting a tubular cell population with an ongoing injury/repair response (Kumar et al. 2015). Increased SOX-9 protein levels were validated in male *Ehhadh* KO renal homogenates (**Fig. S3A**). We also found an increase in Ki67^+^ cells in male *Ehhadh* KO kidneys when compared with WT kidneys (**Fig. 3D**), which is consistent with the RNA-seq data (**Table S1**). Ki67 labels cycling cells (Yang et al. 2010), thus showing an increase in the number of proliferating epithelial cells in *Ehhadh* KO male kidneys. These results show that EHHADH deficiency causes a male-specific PT injury characterized by scattered KIM-1^+^ cells and the detection of regenerating SOX-9^+^ and Ki67^+^ cells.

### Metabolic profiling unveils peroxisomal dysfunction and an increase in glycosphingolipid levels in male Ehhadh KO kidneys

To assess the impact of EHHADH deficiency on kidney metabolism in a non-biased way, we performed metabolite profiling in kidneys from 30-40-week-old WT (n=7) and *Ehhadh* KO (n=5) male mice (**Table S2**). Within the mouse kidneys, 838 known metabolites were detected and quantified. We found 190 metabolites that were significantly altered in the *Ehhadh* KO kidneys compared to WT kidneys (adj p < 0.05), of which 100 metabolites were increased and 90 metabolites were decreased. Increased metabolites in *Ehhadh* KO kidneys include fatty acids and their conjugates, notably those that are markers of peroxisomal dysfunction, such as tetracosahexaenoic acid (C24:6n-3) (Ferdinandusse et al. 2001), pipecolate (Mihalik et al. 1989), and very long-chain acylcarnitines such as C26-carnitine and C26:1-carnitine (Klouwer et al. 2017).

Next, we performed pathway enrichment analysis. We unveiled a significant enrichment of the “Sphingolipid metabolism” pathway among the KEGG pathways (**Fig. 4A**). Indeed, the top seven metabolites increased in *Ehhadh* KO kidneys were glucosylceramides and lactosylceramides, which are all sphingolipids (**Fig. 4B and Table S2**). In the RNA-seq data, we noted an increase in the expression of *Sptlc2* (serine palmitoyltransferase, long chain base subunit 2), the enzyme that initiates *de novo* sphingolipid biosynthesis (Hanada et al. 1997). We also found a male-specific increase in the protein level of SPTLC2 in kidney homogenates from *Ehhadh* KO mice (**Fig. 4C**). Other metabolites that were altered in *Ehhadh* KO mice included increased monoacylglycerols and polyamines, and decreased deoxyribonucleosides, deoxyribonucleotides, glutarylcarnitine and adipoylcarnitine. (**Table S2**). These results suggest that EHHADH deficiency leads to a profound metabolic remodeling in male mouse kidney with a prominent increase in glycosphingolipid levels.

**Figure 4.**
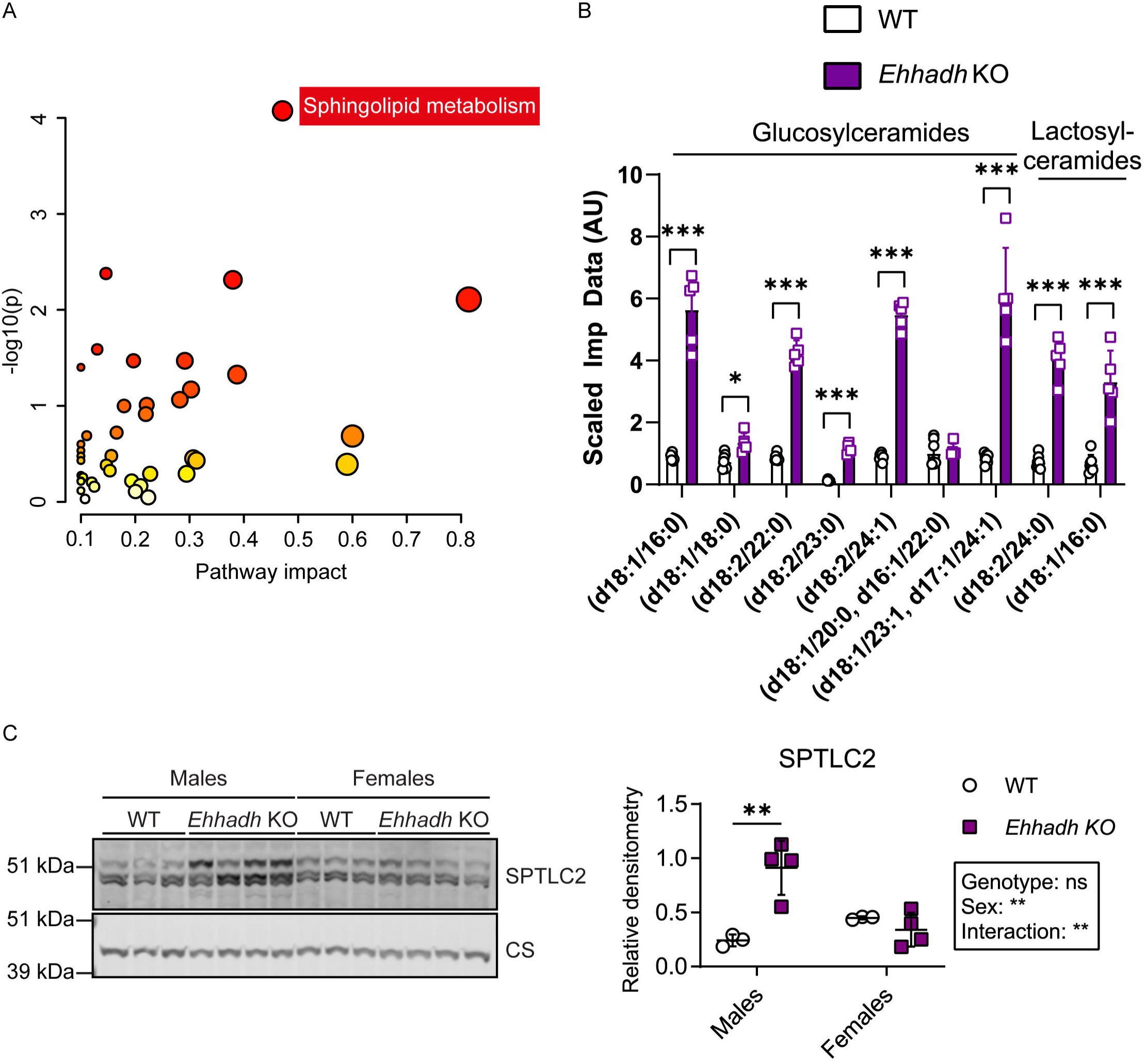
Metabolite profiling in *Ehhadh* KO male kidneys. **A)** Pathway enrichment analysis using significantly altered HMDB metabolites. Scatterplot represents unadj p values from integrated enrichment analysis and impact values from pathway topology analysis. The node color is based on the p values and the node radius represents the pathway impact values. The only significantly altered KEGG pathway using Fisher exact t-test (adj p < 0.05) was “Sphingolipid metabolism”. **B)** Glucosyl- and lactosyl-ceramides levels in WT (n=7) and *Ehhadh* KO (n=5) male mice. The scaled imputed data (Scaled Imp Data) represent the normalized raw area counts of each metabolite rescaled to set the median equal to 1. Any missing values are imputed with the minimum value. **C)** Immunoblots of SPTLC2 and the loading control citrate synthase in WT (n=3 per sex) and *Ehhadh* KO (n=4 per sex) mice, and the corresponding quantification. Data are presented as mean ± SD with individual values plotted (**B, C**). Statistical significance was tested using unpaired t test with Welch’s correction (**B**) or two-way ANOVA with “Genotype” and “Sex” as the two factors, followed by Tukey’s multiple comparisons test (**C**). *P < 0.05; **P < 0.01; ***P < 0.001.

### The kidney phenotype caused by EHHADH deficiency in mice is androgen-dependent

We hypothesized that androgens mediate the male-specific kidney phenotype in *Ehhadh* KO mice. To test this, we first studied the progression of kidney-to-BW ratio in WT and *Ehhadh* KO male mice, from 2 to 52 weeks of age. We found that the kidney-to-BW ratio began to increase after 10 weeks of age, the time when mice reach sexual maturity and plasma testosterone levels peak (Bell 2018). To determine if androgens are directly mediating the kidney phenotype caused by EHHADH deficiency, we compared WT and *Ehhadh* KO mouse kidneys 6 weeks after orchiectomy or a sham operation. Orchiectomy decreased kidney-to-BW ratio in WT and *Ehhadh* KO mice (**Fig. 5B**). Sham operated *Ehhadh* KO mice displayed KIM-1^+^ cells in the cortical area of the kidney, whereas KIM-1^+^ cells were rarely present or even absent in orchiectomized *Ehhadh* KO mice (**Fig. 5C**). We conclude that androgens mediate the kidney enlargement and PT injury in *Ehhadh* KO mice.

**Figure 5.**
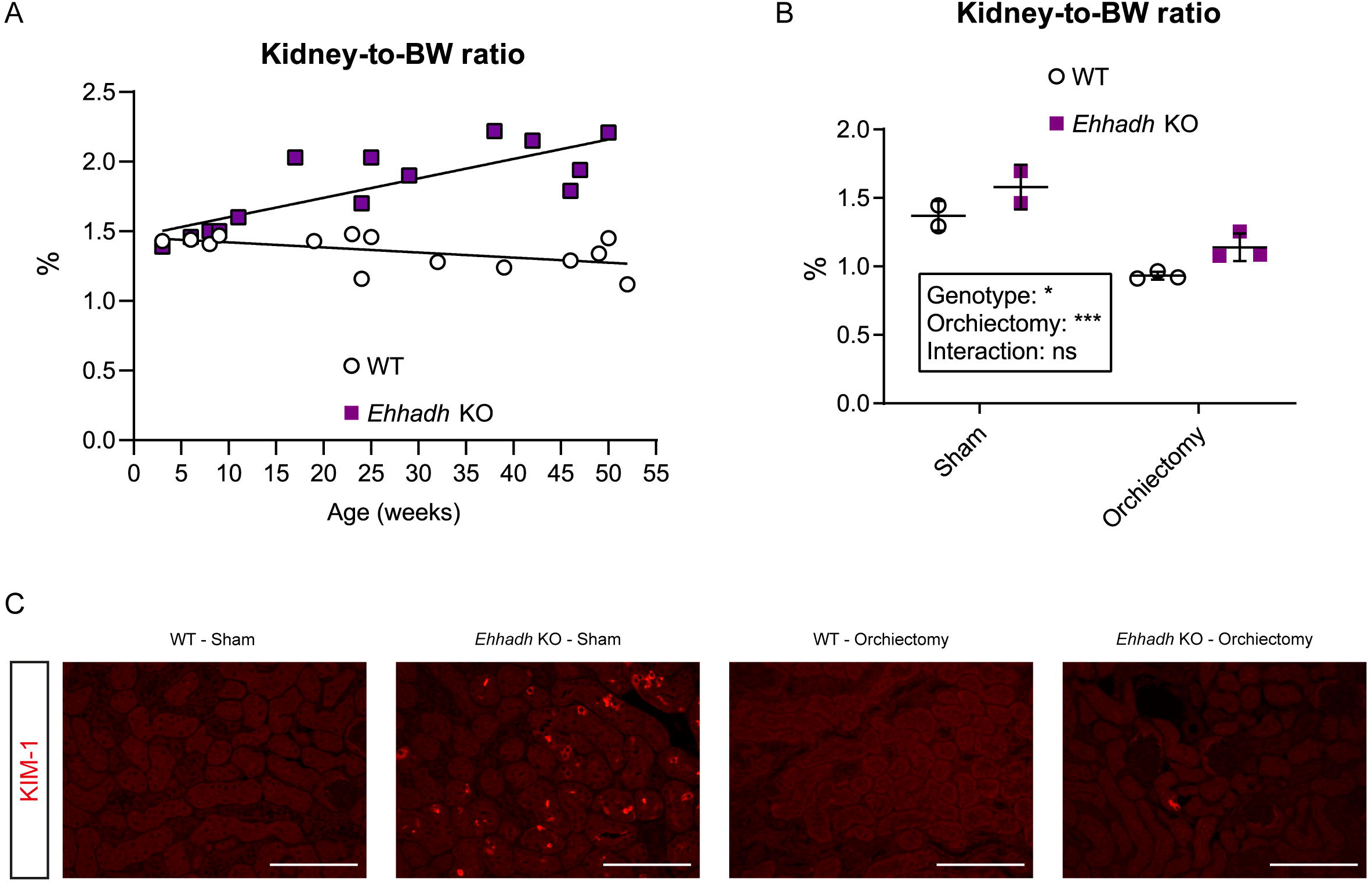
The kidney phenotype caused by EHHADH deficiency in mice is androgen-dependent. **A**) Kidney-to-BW ratio progression with age in male WT and *Ehhadh* KO mice. **B**) Kidney-to-BW ratio in sham-operated (n=2 per genotype) and orchiectomized (n=3 per genotype) WT and *Ehhadh* KO male mice. **C**) Representative images of KIM-1 IF in sham-operated and orchiectomized WT and *Ehhadh* KO male mice. Statistical significance was tested using two-way ANOVA with “Genotype” and “Orchiectomy” as the two factors (**B**). *P < 0.05; ***P < 0.001. Scale bars = 100 µm

### Peroxisomal FAO proteins are sexually dimorphic in mouse kidneys

Several reports have shown significant gene expression differences between sexes in the adult mouse kidney (Ransick et al. 2019; Rinn et al. 2004; Si et al. 2009; Wu et al. 2020). We performed pathway enrichment analysis on two available datasets, one comparing DEGs between male and female healthy kidneys of BALB/c mice (Si et al. 2009), and the other one a PT-specific translational profile comparing male versus female PT cells from mouse kidneys (Wu et al. 2020). We revealed a significant enrichment in genes encoding peroxisomal proteins in both datasets, among other pathways that included fatty acid metabolic pathways (**Fig. S4A, S4B and Table S3**). We then analyzed the levels of peroxisomal FAO proteins in healthy male and female C57BL/6N mouse kidneys. ACOX1 (acyl-CoA oxidase 1), SCPx (sterol carrier protein x, encoded by *Scp2*), CROT (peroxisomal carnitine O-octanoyltransferase) and AMACR (α-methylacyl-CoA racemase) protein levels were higher in male kidneys (**Fig. 6A**). Notably, EHHADH was the only peroxisomal FAO protein whose levels were higher in female kidneys (**Fig. 6A**). No change was found for ABCD3 (ATP-binding cassette sub-family D member 3), DBP (D-bifunctional enzyme) or ACAA1 (3-ketoacyl-CoA thiolase, peroxisomal). To determine if the sexual dimorphism was specific for peroxisomal FAO proteins, we also measured protein levels of two mitochondrial FAO proteins, CPT2 (carnitine palmitoyltransferase 2) and MCAD (medium-chain acyl-CoA dehydrogenase). We found increased levels of MCAD in female kidneys, and no change in the levels of CPT2 (**Fig. 6A**). These results show that most of the peroxisomal FAO proteins show pronounced sexual dimorphism in mouse kidney.

**Figure 6.**
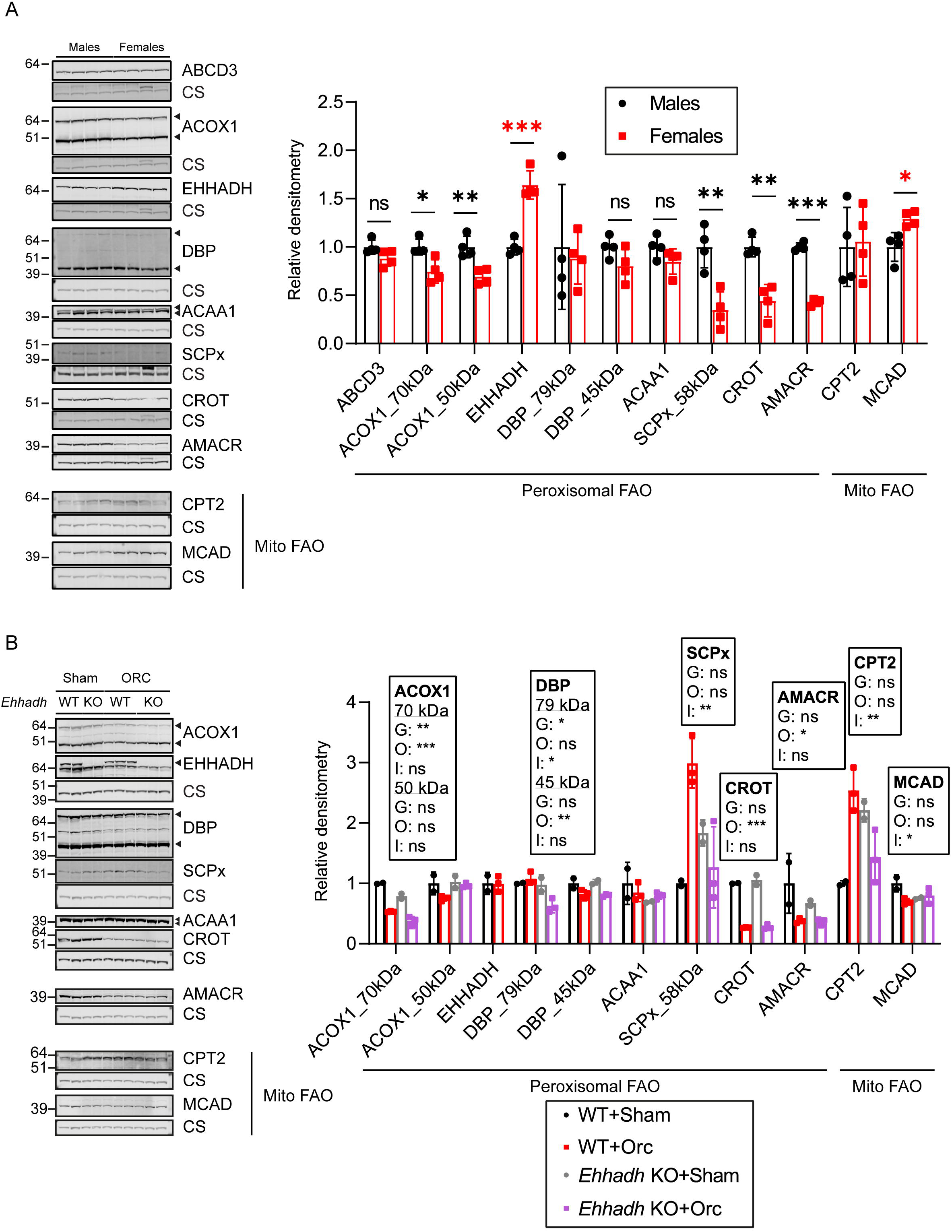
Sexually dimorphic expression of proteins involved in peroxisomal FAO in mouse kidneys. **A**) Immunoblots of peroxisomal and mitochondrial (Mito) FAO proteins with corresponding loading control (citrate synthase) in male and female WT mice (n=4 per sex), and the corresponding quantification. Black asterisks are shown when protein levels were significantly higher in males. Red asterisks are shown when protein levels were significantly higher in females. **B**) Immunoblots of peroxisomal and mitochondrial (Mito) FAO proteins in sham-operated (n=2 per genotype) and orchiectomized (n=3 per genotype) WT and *Ehhadh* KO male mice. Data are presented as mean ± SD with individual values plotted. Statistical significance was tested using unpaired t test with Welch’s correction (**A**) or two-way ANOVA with “Genotype” (G) and “Orchiectomy” (O) as the two factors, (“Interaction”: I) (**B**). *P < 0.05; **P < 0.01; ***P < 0.001.

Next, we investigated if androgens play a role in the regulation of the expression of peroxisomal FAO protein levels in male mouse kidneys. We found that orchiectomy decreased the levels of ACOX1, CROT and AMACR in the kidneys of WT mice (**Fig. 6B**). The levels of these 3 proteins were higher in the kidneys of healthy male WT mice compared with female mice (**Fig. 6A**), suggesting that androgens modulate the levels of these peroxisomal proteins in the kidney. DBP protein levels were also decreased after orchiectomy but only significantly in the kidneys of *Ehhadh* KO mice (**Fig. 6B**). We did not find any effect of orchiectomy in the levels of EHHADH and ACAA1 (**Fig. 6B**). SCPx protein levels showed a differential response to orchiectomy, increasing after orchiectomy in WT kidneys but decreasing in *Ehhadh* KO kidneys (**Fig. 6B**). CPT2 protein levels followed a pattern that was similar to SCPx (**Fig. 6B**), whereas MCAD protein levels tended to decrease after orchiectomy only in WT kidneys (**Fig. 6B**). Altogether, these data provide evidence of a sexual dimorphism in peroxisomal FAO protein content in mouse kidneys.

## Discussion

In 1954, Rhodin discovered peroxisomes in the PT cells of mouse kidneys, and coined them microbodies (Rhodin 1954). And although peroxisomes are highly abundant in the kidney, their function in this organ remains largely unknown. This study establishes a critical function for the peroxisomal L-bifunctional protein (EHHADH) in male renal metabolic homeostasis and highlights the role of peroxisomes in kidney function, in particular in the PT.

The renal phenotype of *Ehhadh* KO mice is associated with a transcriptional signature that shares many features with models of PT injury after AKI (Dhillon et al. 2021; Kirita et al. 2020; Wu et al. 2020), and is characterized by a downregulation of fatty acid metabolic processes and an upregulation of inflammatory pathways. Notably, PT hypertrophy and injury develop spontaneously in *Ehhadh* KO mice in a male-specific fashion. Despite the larger size of male *Ehhadh* KO kidneys, GFR was decreased. Furthermore, the PT of *Ehhadh* KO male mice displayed many KIM-1, SOX-9 and Ki67 positive cells. Therefore, we postulate that the *Ehhadh* KO mouse constitutes a model for male-specific metabolic PT injury. Our data suggest that a reduction in EHHADH activity and peroxisomal FAO may contribute to the development of kidney disease.

In keeping with the notion of a prominent role for peroxisomes in kidney function are observations that peroxisomal abundance and function are decreased in several models of kidney disease, such as PKD (Polycystic kidney disease) (Malas et al. 2017) and AKI models (Gulati et al. 1992; Kalakeche et al. 2011; Kang et al. 2015; Negishi et al. 2007; Ruidera et al. 1988). Patients with peroxisomal disorders such as the Zellweger Spectrum Disorder (ZSD) and DBP deficiency (Huyghe et al. 2006) are prone to develop kidney pathology in the form of renal cysts and/or calcium oxalate stones (Goldfischer et al. 1973; Steinberg et al. 2006; van Woerden et al. 2006). However, underlying mechanisms that link peroxisomal dysfunction with kidney damage other than defective glyoxylate metabolism have not been identified yet. Mouse models with defects in peroxisome biogenesis factors (peroxins) show a more severe renal phenotype when subjected to kidney injury (Maxwell et al. 2003; Weng et al. 2014) . However, peroxin deficiencies impair all peroxisomal functions, hindering the study of the specific role of peroxisomal FAO in kidney physiology. A dominant negative mutation in the *EHHADH* gene was found in patients with an inherited form of renal Fanconi’s syndrome, but the pathophysiological cause was the disruption in mitochondrial oxidative phosphorylation caused by the mistargeting of the mutant EHHADH protein to mitochondria (Assmann et al. 2016; Klootwijk et al. 2014). This is the first study reporting the renal consequences of a single peroxisomal enzyme deficiency in a mouse model.

Our metabolomics analysis provides some insight into the mechanisms underlying the PT injury in *Ehhadh* KO mice. Among the metabolites that accumulated in male *Ehhadh* KO mouse kidneys were very long-chain acylcarnitines (C26- and C26:1-carnitine), pipecolate and tetracosahexaenoic acid (C24:6n-3). Glutarylcarnitine and adipoylcarnitine were decreased. The direction of change in these metabolites is consistent with defective peroxisomal metabolism (Ferdinandusse et al. 2001; Klouwer et al. 2017; Mihalik et al. 1989). Our data suggest that EHHADH deficiency induces a rewiring of cellular metabolism that results in the accumulation of glucosyl- and lactosylceramides in the kidney, with many of these species containing at least one very long-chain acyl chain (C22 or longer). The accumulation of these complex sphingolipids has been associated with an increase in kidney size mediated by androgens (Coles et al. 1970; Hay & Gray 1970; Koenig et al. 1980; McCluer et al. 1981), but a link with peroxisomes has not been established.

The primary metabolic cause of the male-specific renal phenotype of *Ehhadh* KO mice remains unknown. We speculate that EHHADH deficiency impairs the degradation of a metabolite that is toxic to the PT epithelium and envision two possible models to explain this phenomenon (**Fig. S5**). In the first model (A), such a toxic metabolite is produced by one of the CYP enzymes whose expression is known to be sexually dimorphic (Rinn et al. 2004; Si et al. 2009; Wu et al. 2020). Dicarboxylic acids and eicosanoids are likely candidate metabolites as both undergo several cycles of peroxisomal β-oxidation after initial ω-oxidation by CYP enzymes (de Waart et al. 1994; Diczfalusy 1994; Diczfalusy et al. 1991; Dirkx et al. 2007; Ferdinandusse et al. 2002, 2004; Gordon et al. 1990; Mayatepek et al. 1993; Ranea-Robles et al. 2021; Tomoko et al. 1986). Unfortunately, our metabolomics analysis did not reveal a clear candidate toxic molecule, but the detection may have been obscured by the ongoing disease process. Future studies should focus on differences between the metabolome of male and female kidneys focusing on lipid substrates of CYP enzymes.

In the alternative model (B), the toxic insult caused by EHHADH deficiency may occur in both males and females, but only progresses to PT injury in males due to higher androgen levels. We found that the relative kidney enlargement was first detectable around 10 weeks of age in male *Ehhadh* KO mice, right after plasma androgen levels peak in mice (Bell 2018). Moreover, castration reversed kidney enlargement and PT injury in male *Ehhadh* KO mice. We also found a male-specific increase in SPTLC2 protein levels in renal homogenates. SPTLC2 catalyzes the first and rate-limiting step of sphingolipid biosynthesis (Hanada et al. 1997), suggesting that the metabolic rewiring caused by EHHADH deficiency is also restricted to male animals.

Sex-specific differential susceptibility is reproduced in other rodent models of kidney disease (Müller et al. 2002; Park et al. 2004). Androgens and, in particular, testosterone, are known to induce renal hypertrophy in rodents (Pfeiffer et al. 1940; Selye 1939) and to increase the susceptibility of mice to develop kidney injury (Park et al. 2004). In humans, the available evidence suggests that male gender is associated with a more rapid rate of disease progression and a worse renal outcome in patients with CKD (Neugarten & Golestaneh 2019; Neugarten et al. 2000). Moreover, a recent study found that genetically predicted levels of testosterone in men were associated with higher risk of CKD and worse kidney function (Zhao & Schooling 2020).

In conclusion, we show that deficiency of a single peroxisomal FAO enzyme, EHHADH, causes kidney hypertrophy, GFR decrease and PT injury in male mice. The results of our study suggest that EHHADH and peroxisomal metabolism play an important role in metabolic homeostasis of the PT. Altogether, this study underlines the role of peroxisomes and EHHADH in the mouse kidney and provides evidence for a sexual dimorphic pathophysiologic mechanism within the kidney.

### Author contributions

P.R-R., K.L., J.C.H, and S.M.H. designed the study; P.R-R., A.B., K.P., K.L., and D.M. carried out experiments; P.R-R., A.B., C.A., and S.M.H. analyzed the data; P.R-R., C.A., and S.M.H made the figures; P.R-R. and S.M.H drafted the original version of the manuscript; all authors revised the paper and approved the final version of the manuscript

## Acknowledgements

We thank Dr. Sacha Ferdinandusse for providing the AMACR antibodies, and the Mount Sinai School of Medicine Biorepository and Pathology Core for their help with histology preparations.

Research reported in this publication was supported by the National Institute of Diabetes and Digestive and Kidney Diseases of the National Institutes of Health under Award Number R01DK113172. The content is solely the responsibility of the authors and does not necessarily represent the official views of the National Institutes of Health.

