## Supplementary figures and images for "Deficiency of peroxisomal L-bifunctional protein (EHHADH) causes male-specific kidney hypertrophy and proximal tubular injury in mice"

### Supplementary Figure S1

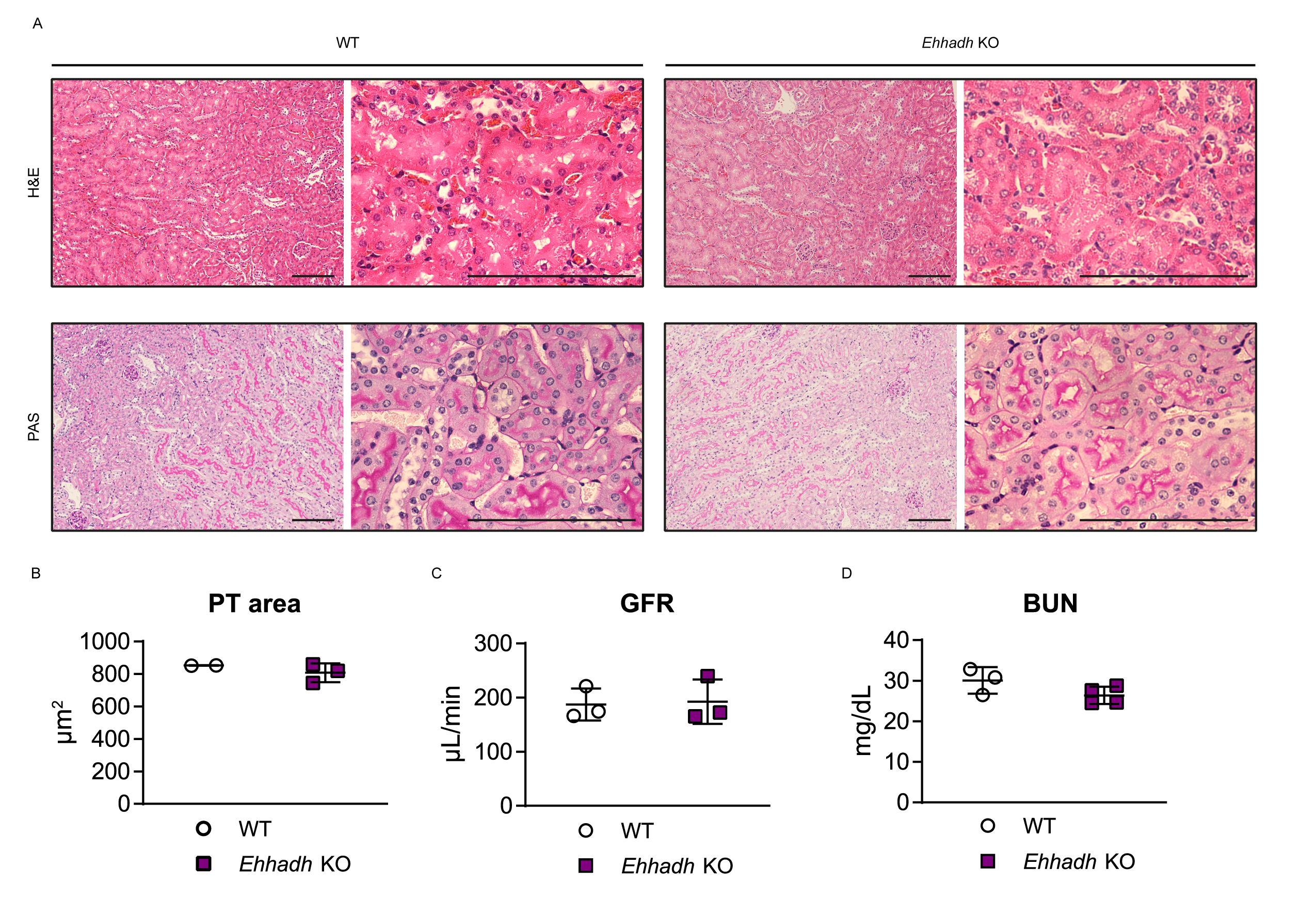

### Supplementary Figure S2

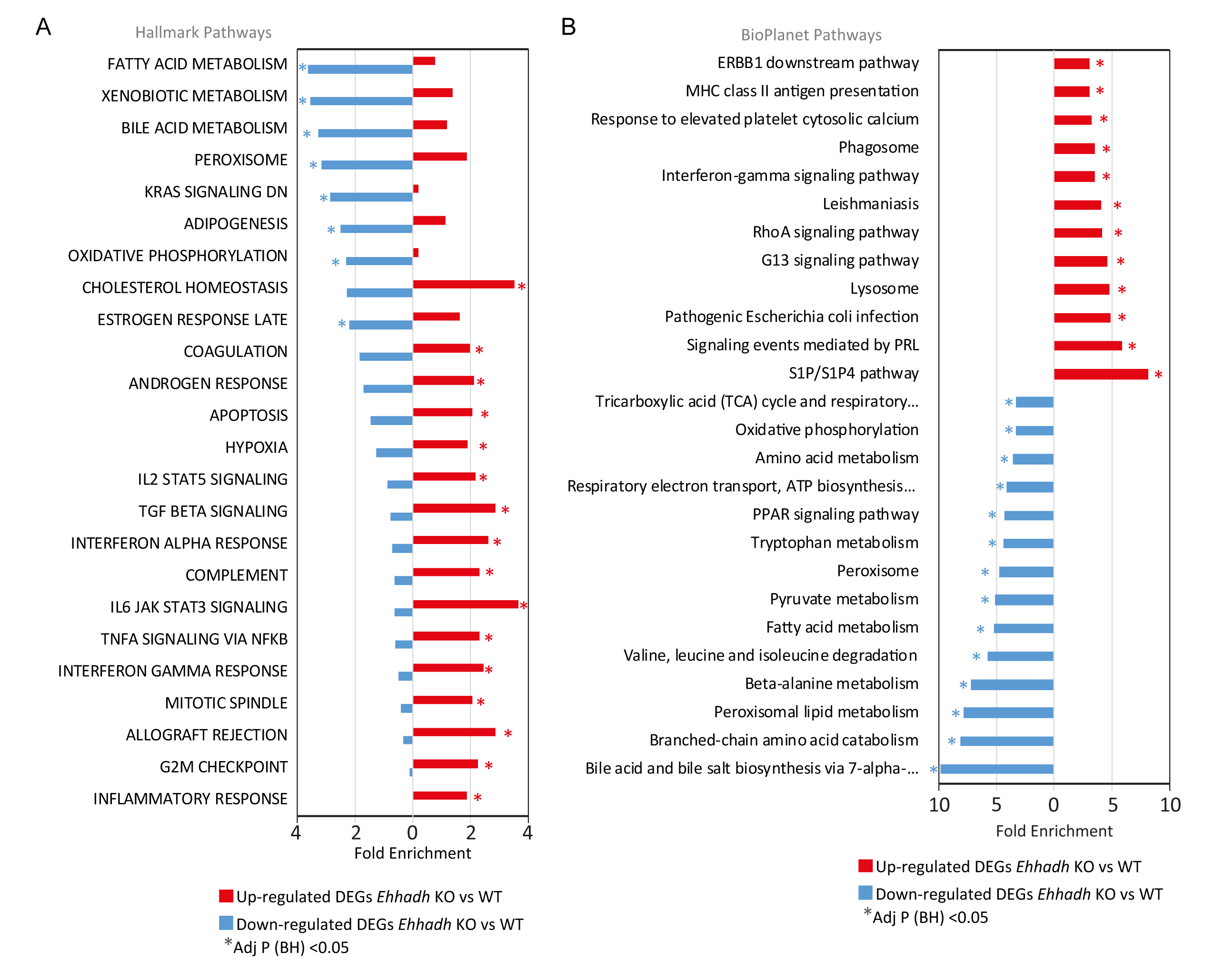

### Supplementary Figure S3

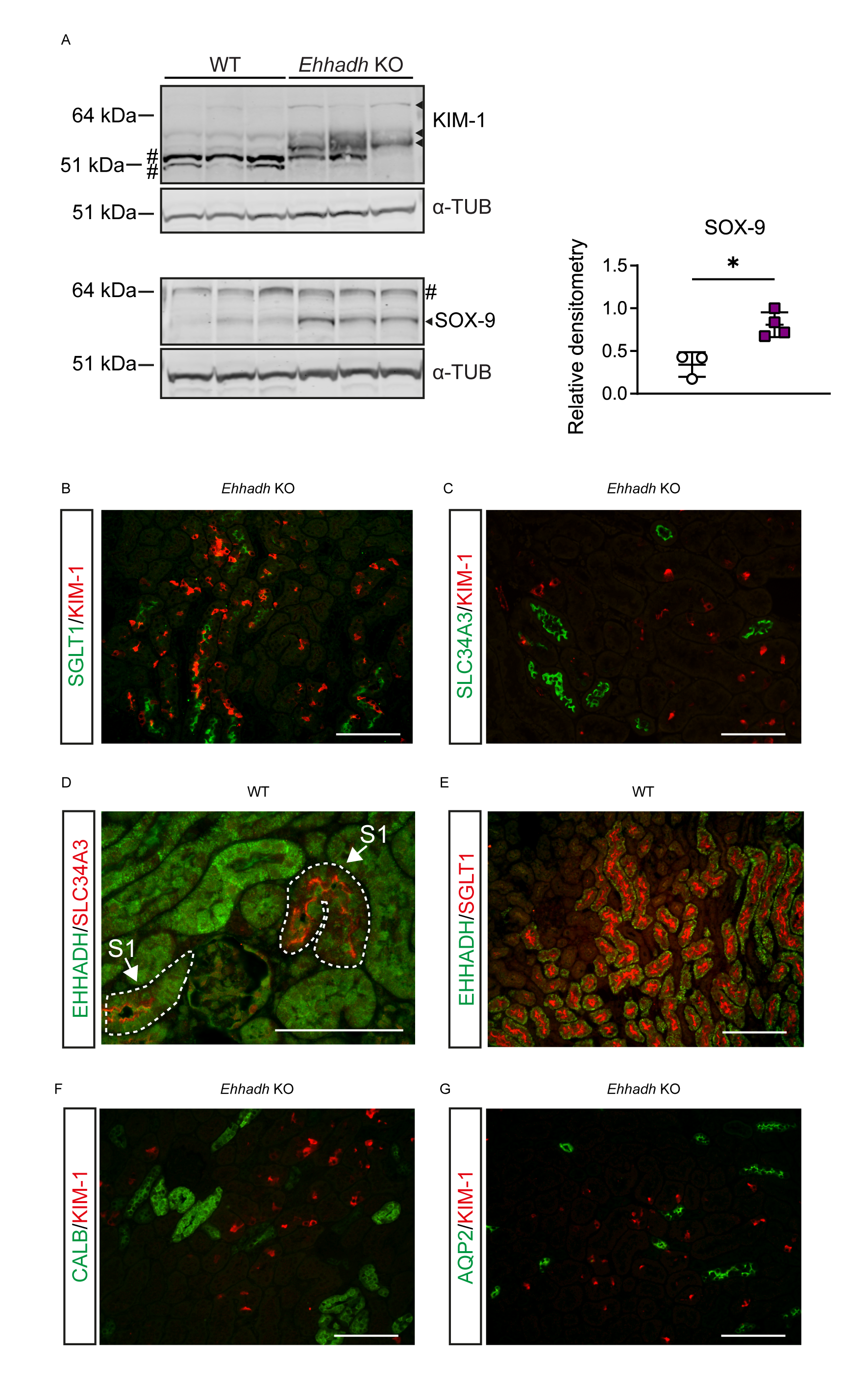

### Supplementary Figure S4

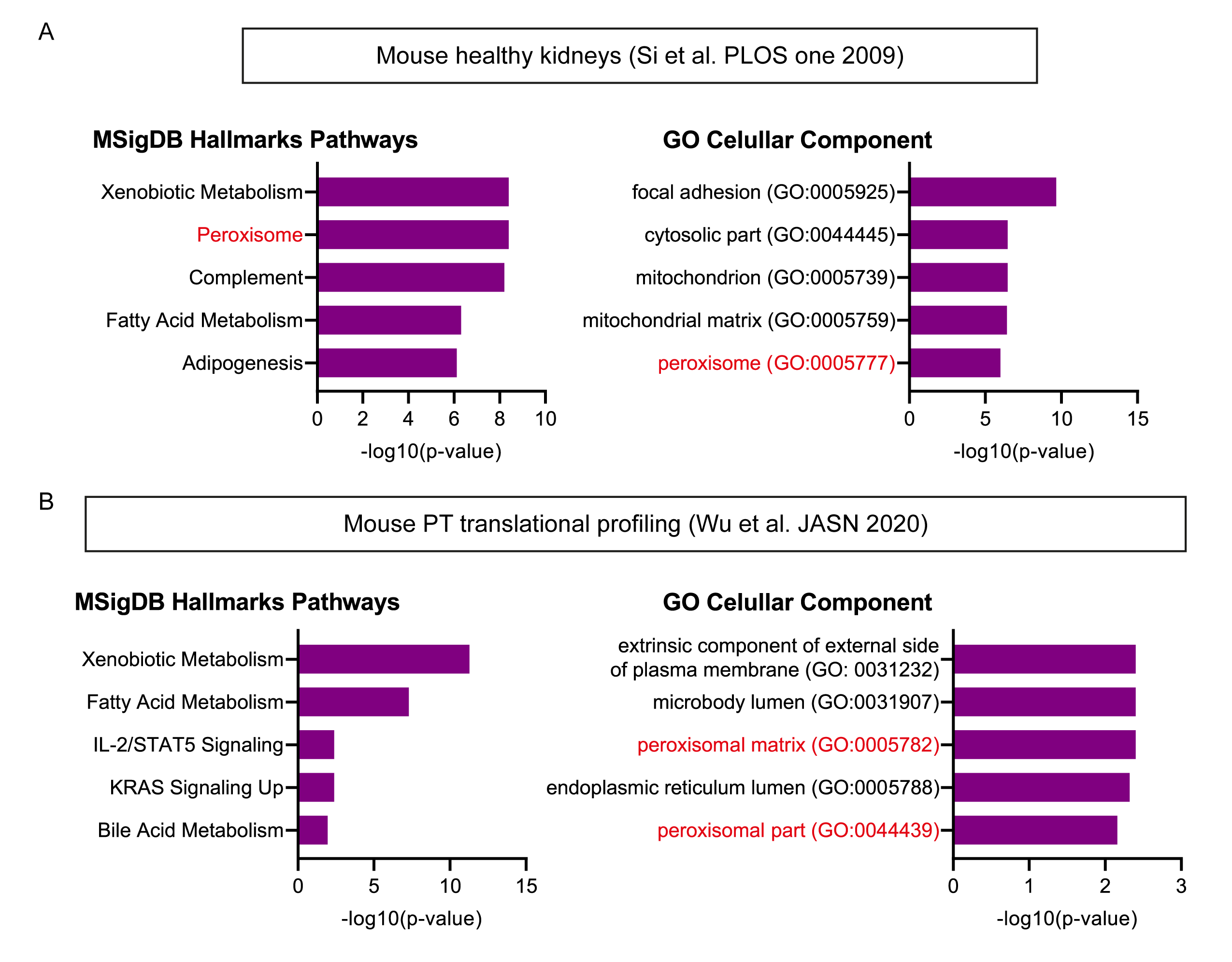

### Supplementary Figure S5

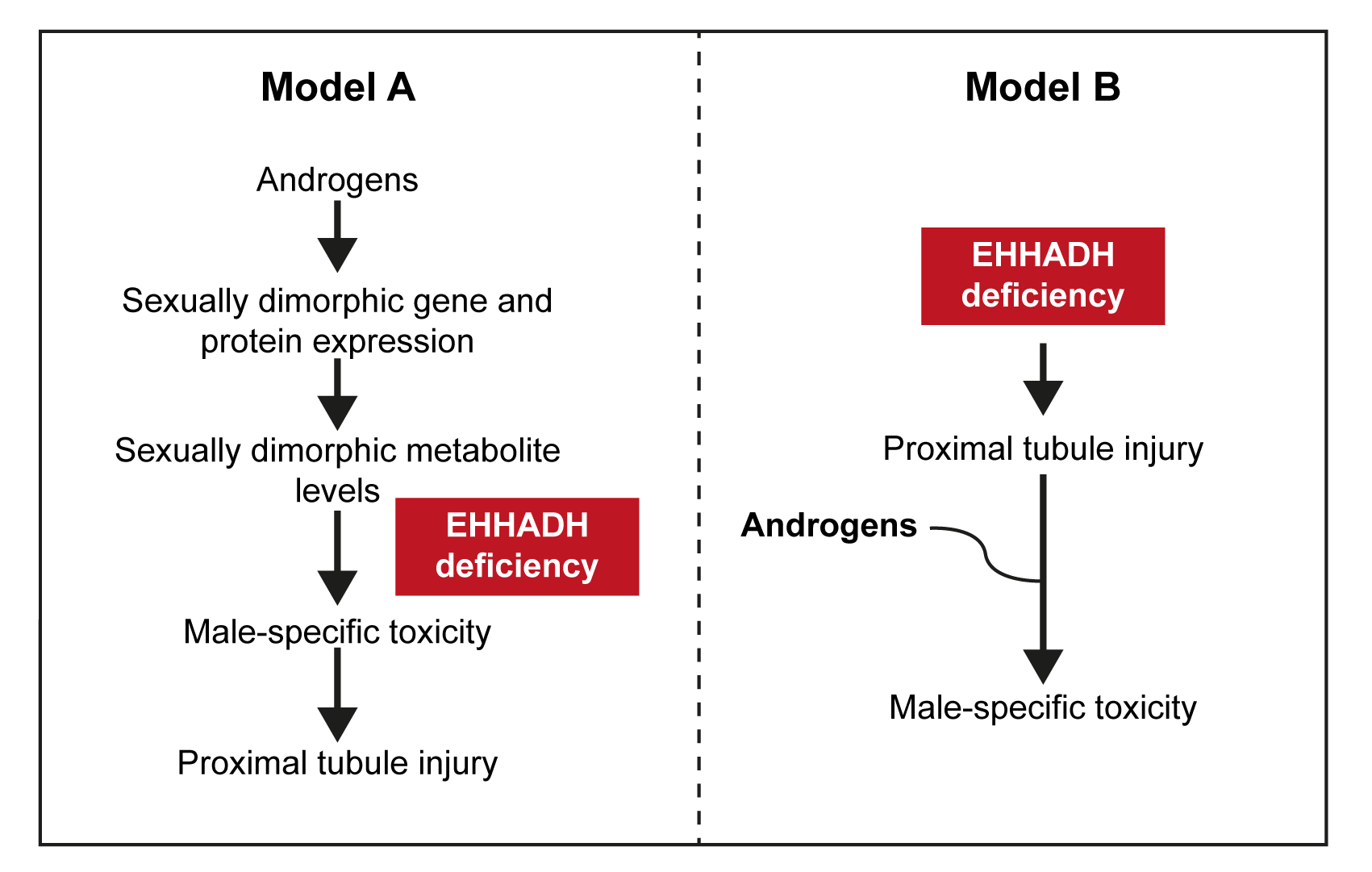
