## Supplemental information for "Deficiency of peroxisomal L-bifunctional protein (EHHADH) causes male-specific kidney hypertrophy and proximal tubular injury in mice"

**Table of Contents**

**Supplementary Figures**

**Supplementary Figure S1.** EHHADH deficiency does not cause morphological or functional changes in female mouse kidneys

**Supplementary Figure S2.** Transcriptomics of male Ehhadh KO kidneys

**Supplementary Figure S3.** EHHADH deficiency activates the proximal tubule injury response in male mice

**Supplementary Figure S4.** Pathway enrichment analysis of sexually dimorphic DEGs in mouse kidneys

**Supplementary Figure S5**. Proposed working models to explain how EHHADH deficiency causes male-specific PT injury

**Supplementary Tables**

**Supplementary Table S1.** *Ehhadh* KO kidney RNA-seq signature

**Supplementary Table S2.** Untargeted metabolomics dataset comparing WT and *Ehhadh* KO male kidneys.

**Supplementary Table S3.** Pathway enrichment analysis of two published datasets containing sexually dimorphic genes in mouse kidneys

**Supplementary Figure S1**. **EHHADH deficiency does not cause morphological or functional changes in female mouse kidneys.**

**A)** Representative images of H&E and PAS staining in kidney sections from female WT and *Ehhadh* KO mice. Scale bars = 100 µm. **B**) Morphometric analysis of the cross-sectional tubule areas in WT (n=2) and *Ehhadh* KO (n=3) female mice. **C**) Glomerular filtration rate (GFR) in WT and *Ehhadh* KO female mice (n=3 per genotype). **D**) Blood urea nitrogen (BUN) levels (mg/dL) in WT (n=3) and *Ehhadh* KO (n=4) female mice. Data are presented as mean ± SD with individual values plotted.

**Supplementary Figure S2**. **Transcriptomics of male *Ehhadh*** **KO kidneys**

Pathway enrichment analysis of genes either significantly up- or down- regulated in *Ehhadh* KO mice versus WT according to Hallmark (**A**) or BioPlanet (**B**) databases. Values represent the fold enrichment and significance is indicated as * adj p<0.05. All BioPlanet pathways shown are significantly enriched as adj p <0.05 and only pathways with fold enrichment >3.0 are shown. Full table of results are in Table S1B and S1C.

**Supplementary Figure S3. EHHADH deficiency activates the proximal tubule injury response in male mice.**

**A**) Immunoblots of KIM-1 and SOX-9, and their loading control α-Tub (alpha-tubulin). SOX-9 protein levels were quantified relative to α-Tub. Several bands showed in the KIM-1 blot correspond to non-specific bands (#), as reported elsewhere (Ichimura et al. 2004). This is in line with the absence of KIM-1 signal in IF studies in WT kidneys. The quantified band for SOX-9 is marked with a black arrowhead. **B)** Representative image of SGLT1 (red) and EHHADH (green) co-immunostaining in the cortical area of a WT mouse kidney. **C**) Representative image of SLC34A3 (red) and EHHADH (green) co-immunostaining in the cortical area of a WT mouse kidney. Two S1 segments are marked. **D**) Representative image of SGLT1 (green) and KIM-1 (red) co-immunostaining in the cortical area of an *Ehhadh* KO male kidney. **E**) Representative image of SLC34A3 (green) and KIM-1 (red) co-immunostaining in the cortical area of an *Ehhadh* KO male kidney. **F**) Representative image of CALB (green) and KIM-1 (red) co-immunostaining in an *Ehhadh* KO male kidney. **G**) Representative image of AQP2 (green) and KIM-1 (red) co-immunostaining in an *Ehhadh* KO male kidney. Data are presented as mean ± SD with individual values plotted. Statistical significance was tested using unpaired t test with Welch’s after multiple comparison correction *P < 0.05. Scale bar = 100 µm.

**Supplementary Figure S4. Pathway enrichment analysis of sexually dimorphic DEGs in mouse kidneys**

**A**) Top 5 significant MSigDB Hallmarks terms and GO Cellular Component terms after pathway enrichment analysis using DEGs from Si et al (Si et al. 2009). B) Top 5 significant MSigDB Hallmarks terms and GO Cellular Component terms after pathway enrichment analysis using DEGs from Wu et al (Wu et al. 2020). Terms were ranked by -log10(adj p).

**Supplementary Figure S5**. **Proposed working models to explain how EHHADH deficiency causes male-specific PT injury**
